# How an arthropod-borne virus manages host switches: exploration of host-specific mutations and gene formula in bluetongue virus

**DOI:** 10.64898/2025.12.04.692277

**Authors:** Yannis Moreau, Patricia Gil, Antoni Exbrayat, Ignace Rakotoarivony, Emmanuel Bréard, Corinne Sailleau, Cyril Viarouge, Stephan Zientara, Giovanni Savini, Maria Goffredo, Giuseppe Mancini, Etienne Loire, Serafin Gutierrez

## Abstract

Arthropod-borne viruses, also known as arboviruses, cause disease worldwide. These viruses alternate between vertebrates and arthropods, facing distinct selection pressures in each host type. Surprisingly, whether specific mechanisms facilitate host alternation remains unclear. Here, we use bluetongue virus (BTV) as a model to investigate two phenomena potentially facilitating host alternation: adaptive mutations and genome formula variation (GFV). First, we searched for mutations indicative of host adaptation in cattle, sheep, and midges infected during an epizooty. We found a limited number of parallel mutations. This result suggests that adaptive mutations are not a main mechanism facilitating host alternation in BTV. Second, we investigated a link of GFV with a proxy of viral fitness, the speed of virus dissemination in midges. We analyzed the genome formula (GF) in experimentally-infected *Culicoides imicola* midges, either with an infection disseminated into the head or not. GFs differed between dissemination levels, suggesting a link between GF and the speed of virus dissemination. Finally, we investigated the mechanisms underlying GF generation. Currently, two scenarios have been proposed: random generation followed by selection, or directed generation through molecular mechanisms. To limit the influence of selection, we analyzed the GF after a single replication cycle in cell culture. GF values converged towards specific values among replicates. Moreover, GFs differed between arthropod and vertebrate cells. Thus, GF generation is not purely stochastic and is probably influenced by host-specific mechanisms. Overall, our findings highlight GFV as a potential mechanism facilitating BTV alternation across hosts, but not classical mutation-driven evolution.

**Author Summary:** Arthropod-borne viruses are important public health problems in the world. These viruses follow a complex cycle involving alternation between arthropods and vertebrates. Surprisingly, we know little on the mechanisms that facilitate alternation between the two host types. Knowledge on such mechanisms could allow the development of antiviral tools. Here, using Bluetongue virus as a model, we investigated two potential mechanisms facilitating alternation: adaptive mutations and genome formula variation (GFV). We found limited evidence for host-specific mutations in natural populations. We observed a potential link between GFV and viral fitness. Moreover, our results suggest that molecular mechanisms generate GFV in a host-specific way from the first round of cell infection.

## Introduction

Arthropod-borne viruses, or arboviruses, are paradigmatic examples of complex viral cycles. These cycles involve switches between vertebrates and arthropods at each transmission event [1]. Several arboviruses are the subject of intense research due to their health impact worldwide [2]. However, the potential mechanisms that facilitate host alternation remain a largely unstudied aspect [3]. This situation is surprising as a deep understanding of host alternation could lead to advances in antiviral design, among other possibilities.

Arboviruses must cope with different selection pressures in each host type. For example, RNA silencing is the main antiviral mechanism in arthropods whereas the adaptive immune system plays this role in vertebrates. Alternating between hosts with different selection pressures should incur a fitness cost, mainly due to antagonistic pleiotropy [4]. Experimental studies with different arboviruses support this prediction [5–8], although counterexamples exist [9–11]. Nevertheless, the arbovirus cycle should also confer benefits as it has evolved among a wide diversity of viruses. Hence, if there are costs associated with their cycle, certain arboviruses may have acquired mechanisms that allow phenotype optimization during host alternation.

Two main phenomena, not mutually exclusive, could allow for this optimization. The first is host adaptation through adaptive mutations. RNA viruses seem particularly well-suited for this process given their high mutation and replication rates [12]. Many studies have explored this phenomenon through experimental infections [5–9,13]. Most of these studies show that specific mutations arise when viral populations are passaged in a single cell type. However, host-specific mutations have rarely been observed in the few studies that have analyzed natural populations (*i.e.*, field samples) [14–17]. Moreover, most of these studies have focused on only a few arboviruses, primarily flaviviruses. This paucity of data leaves unanswered the question of the role of adaptive mutations in host alternation.

The second process allowing phenotype optimization is phenotypic plasticity. Phenotypic plasticity is defined as the ability to produce different phenotypes when exposed to different environmental conditions [18]. Mechanisms leading to phenotypic plasticity have been observed in arboviruses through multifunctional proteins, like the NS1 protein in dengue virus [19,20], or through subgenomic RNAs in flaviviruses [21]. However, phenotypic plasticity has rarely been explored as a mechanism facilitating host alternation in arboviruses [3].

We have previously identified a phenomenon that could allow for phenotypic plasticity and, thus, facilitate host alternation in an arbovirus [3]. This arbovirus is bluetongue virus (BTV; *Orbivirus*, *Reoviridae*), a major pathogen of domestic and wild ruminants worldwide. The BTV cycle involves a compulsory alternation between ruminants and *Culicoides* biting midges (Diptera). BTV has a segmented genome consisting of ten segments of double-stranded RNA. All but one of the segments encode for a single open reading frame (ORF) [22]. BTV particles are thought to contain one copy of each segment [23,24]. This conservative encapsidation should result in an equal number of copies of each segment in BTV populations. However, the BTV genome segments are not present in equimolar amounts in field populations infecting ruminants and biting midges [3]. Interestingly, differences in copy number between segments are specific to each host type. These observations are reminiscent of a phenomenon called genome-formula variation (GFV) in plant viruses [25,26]. GFV is thought to allow for phenotypic plasticity.

The genome formula (GF) is a parameter consisting of the relative frequencies of all viral segments in a within-host population. GFV takes place when distinct and reproducible GFs occur among hosts. These host-specific GFs are known as set-point GFs and could allow for phenotype optimization in different hosts [25,27,28]. Under this hypothesis, differences in gene frequencies specifically regulate gene expression in each host, thereby optimizing the phenotype. Thus, GFV resembles the well-known phenomenon of variation in gene copy number in eukaryotes and bacteria [29]. However, GFV operates at the level of the virus population rather than at the level of the individual genome. GFV was first identified in multipartite viruses of plants (*i.e.* viruses with a segmented genome where each segment is encapsidated separately) [25]. GFV-like phenomena have also been observed in three unrelated segmented viruses: BTV, influenza A virus (*Orthomyxoviridae*) and tomato spotted wilt virus (*Tospoviridae*) [3,30,31]. These observations suggest that GFV may be more widespread than previously thought. Nevertheless, GFV remains largely unstudied in viruses other than multipartite ones. For example, little is known about its potential influence on viral fitness or the mechanisms that generate GFs.

Here, we address three key questions regarding host alternation in BTV. First, we investigate the potential involvement of adaptive mutations in host alternation. To this end, we have analyzed the BTV genetics in natural infections of vertebrates and insects. Previous studies have shown that these populations exhibit GFV-like behavior [3]. Second, we study the potential influence of GFV on viral fitness through experimental infections of biting midges. Finally, we examine whether random processes lead to GFV using cell culture infections.

## Results

### Scarcity of parallel mutations suggestive of host adaptation

We examined the genetics of wild BTV populations to determine if they suggest host adaptation. To this end, we sequenced the genomes of BTV populations from natural infections in cows, sheep and *Culicoides* biting midges from the same epizooty. These populations bear GFs specific to each host [3]. We searched for SNPs specific to a given host and present in several samples of that host. Such parallel mutations strongly suggest convergent evolution during host adaptation.

To avoid mutations associated with passages in cell culture, we developed a sequencing protocol that allows for direct sequencing of field samples. The protocol was used to sequence 21 samples from cows, sheep and biting midges. Sequencing generated 5.22 × 10^9^ bases per sample (20.8 million reads per sample). Thirteen samples were selected for SNP analysis after sample selection based on coverage (five midge, four cow and four sheep samples). Out of 18 571 genome positions, 70 positions contained a SNP with a derived allele frequency above 0.2 (Figure 1, Table S1). No site was three-allelic.

**Figure 1.**
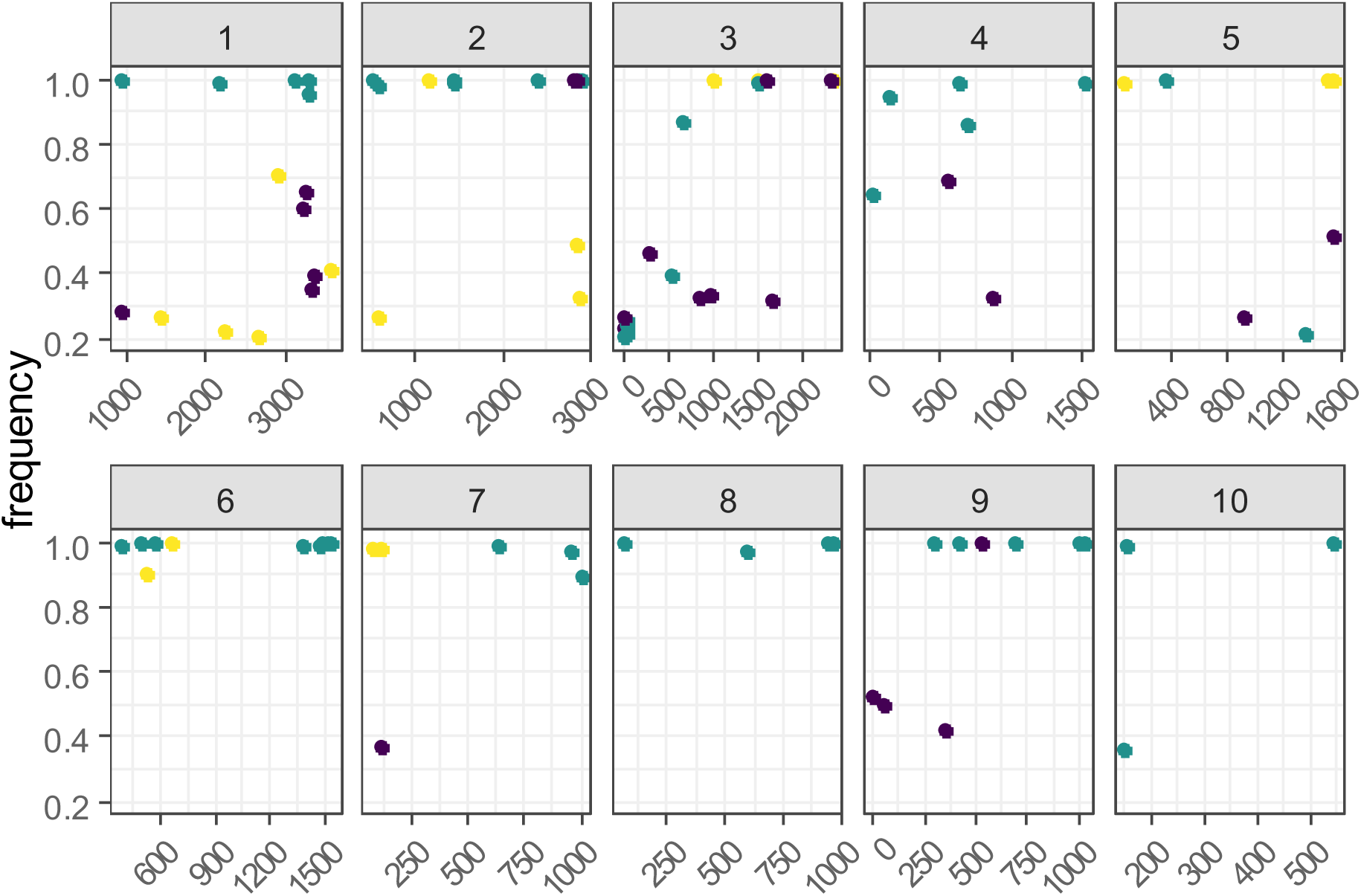
Single nucleotide polymorphisms (SNPs) along the genome of Bluetongue virus and among hosts. Each facet corresponds to a segment and the segment number appears at the top of each facet. The x-axis indicates segment lengths and the y-axis indicates allele frequency. Dots indicate SNPs and their color stands for the host in which the SNP was observed (black: cow, green: midges, yellow: sheep).

First, we searched for fixed SNPs shared by several samples from the same host. Such parallel mutations are a classical signature of host adaptation. We found 45 SNPs (64%) fixed or close to fixation. Fixed SNPs were detected in all segments and hosts, with a higher number of fixed SNPs in midge samples (Figure 1). However, no fixed SNP was shared by all samples of any host (Figure 2). Twelve fixed SNPs were found in two samples of the same host (Figure 2, Table S2). These SNPs were distributed over all segments but segments 1 and 10 (Figure 2, Table S2). Most of those SNPs were found in midges (seven out of 12 SNPs; Figure 2, Table S2). Eleven out of the 12 positions were located in coding regions and three SNPs led to non-synonymous changes (Table S2). The non-synonymous mutations were only found in vertebrate hosts (Table S2). We then searched for fixed SNPs common to the two vertebrate hosts, which could indicate adaptation to vertebrates. One such SNP, a non-synonymous SNP, was found in two sheep and one cow (SNP in position 2322 of segment 3; Figure 2A, Table S1). Finally, we searched for SNPs shared by both vertebrates and midges. One fixed synonymous SNP was shared by a sheep sample and a midge sample (SNP in position 514 of segment 6, Table S1).

**Figure 2.**
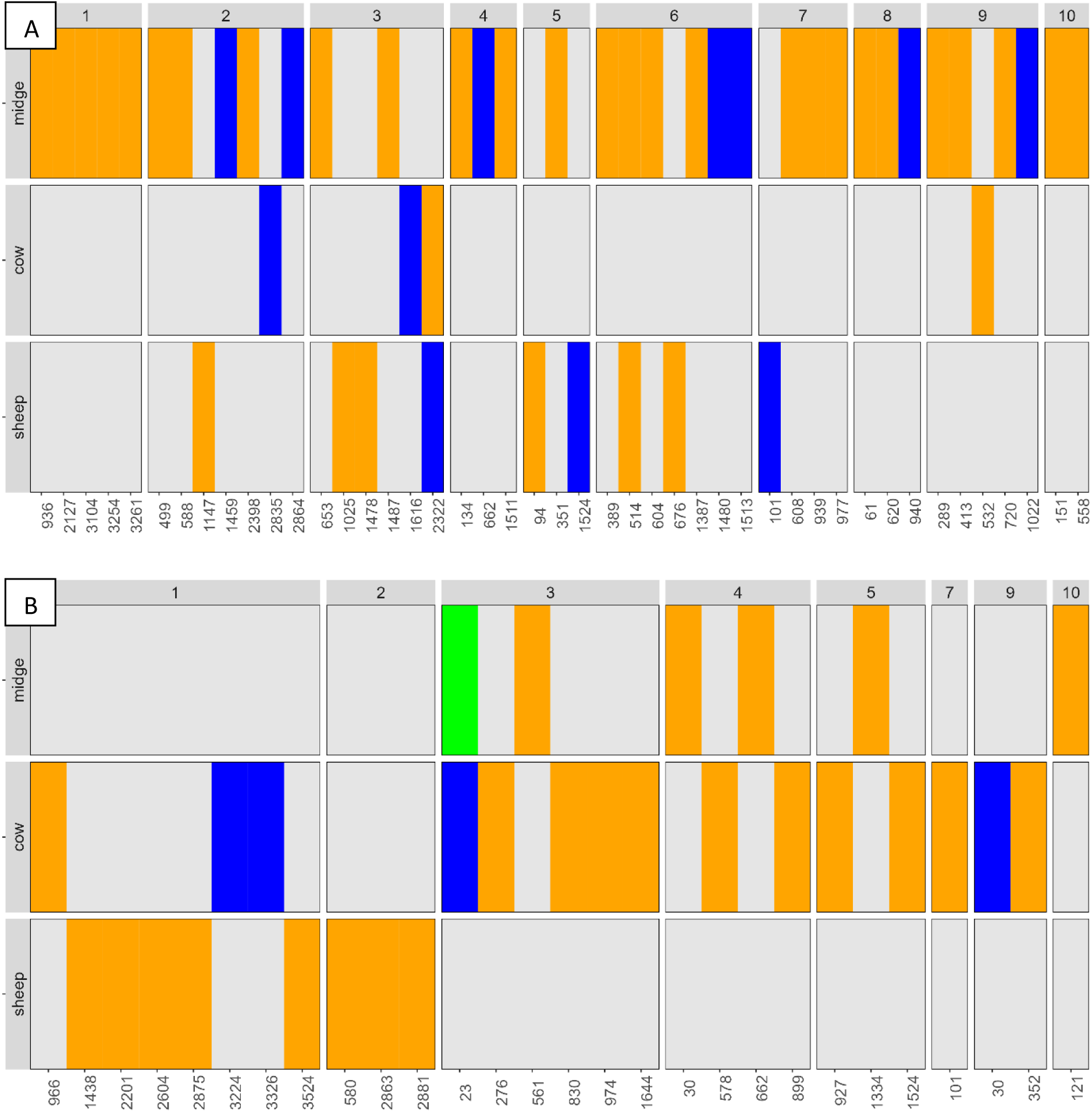
Distribution of SNPs, fixed (A) or non-fixed (B), among segments and hosts. Segment names appear on top. Host names appear on the left. SNP position in each segment is indicated on x-axis. Each SNP is indicated with a colored tile. Tile color stands for the number of samples in which the SNP was found (orange: one sample, blue: two samples, green: four samples).

Certain host-specific SNPs may not have reached fixation at sampling because we could not control for inoculation time in the natural populations. Thus, we searched for non-fixed SNPs shared by several samples of the same host. A total of 28 non-fixed SNPs were found. These SNPs were detected in all hosts, and in all segments except segments 6 and 8 (Figure 2B, Table S1). We detected non-fixed SNPs shared by several samples of the same host only in cows. These fours SNPs were detected in two cow samples (Figure 2B, Table S3). One out of the four non-fixed parallel SNPs led to a non-synonymous mutation (shared by two cows; Table S3). Moreover, we did not observe any position with both fixed and non-fixed SNPs in any host. We only observed positions with both fixed and non-fixed SNPs when analyzing together all vertebrate samples (two SNPs fixed in two sheep and non-fixed in a cow; segment:position = 5:1524 and 7:101; Figure 1, Table S1). Finally, one non-fixed SNP was shared by two cows and four midge samples (position 23 in segment 3, Figure 1, Table S3).

### The speed of within-host dissemination correlates with the genome formula

We analyzed the potential correlation between the GF and a proxy of viral fitness. This proxy is the speed at which the virus disseminates into the head in midges. Head infection is often used as a proxy for salivary gland infection and, thus, probability of transmission to vertebrates [32–34]. Furthermore, the first virions to infect the salivary glands could block subsequent infections through superinfection exclusion [35,36]. Thus, we hypothesized that BTV populations with optimal GFs may disseminate faster within the midge body. We tested this hypothesis in experimental infections of wild *Culicoides imicola* midges [34].

We defined two levels of virus dissemination in midges at ten days after intake of a virus-spiked blood meal. The two levels were a detectable dissemination into the head or no detection. These levels were named full and limited dissemination, respectively. We screened the decapitated bodies and heads of individual midges separately to identify those belonging to each dissemination group. Then, we quantified each of the ten segments in the decapitated bodies. Viral loads in heads were too low to allow for a robust quantification of segment copies (i.e., less than 1000 copies) and were not included in the analysis of segment frequencies. The final dataset consisted of 17 midges with full disseminations and seven midges with limited dissemination.

The distributions of segment frequencies followed a genome-formula behavior (Fig. 3). That is, frequencies significantly differed from equimolarity and between segments (equimolarity test: Wilcoxon signed rank test, Benjamini-Hochberg correction, p-values < 2×10^-4^ except for segment 6, Tab. S4; between-segment test for each dissemination level: ANOVA, p-values < 10^-15^, see Tab. S5 for p-values of pairwise comparisons of segments adjusted with Tukey honest significance test; Fig. 3). Moreover, segment frequencies significantly differed between the two dissemination groups for five segments (general linear model, p-value adjustment with rotation testing, p-values < 0.030 for segments 1, 2, 3, 7 and 8; Fig. 3 and Tab. S6). These differences involved a significantly higher deviation from equimolarity of virus populations with a full dissemination (Wilcoxon rank sum test, p-value = 3×10^-4^, Fig. S1). Hence, a full dissemination correlated with a specific GF, and this GF had a higher divergence from equimolarity.

**Figure 3.**
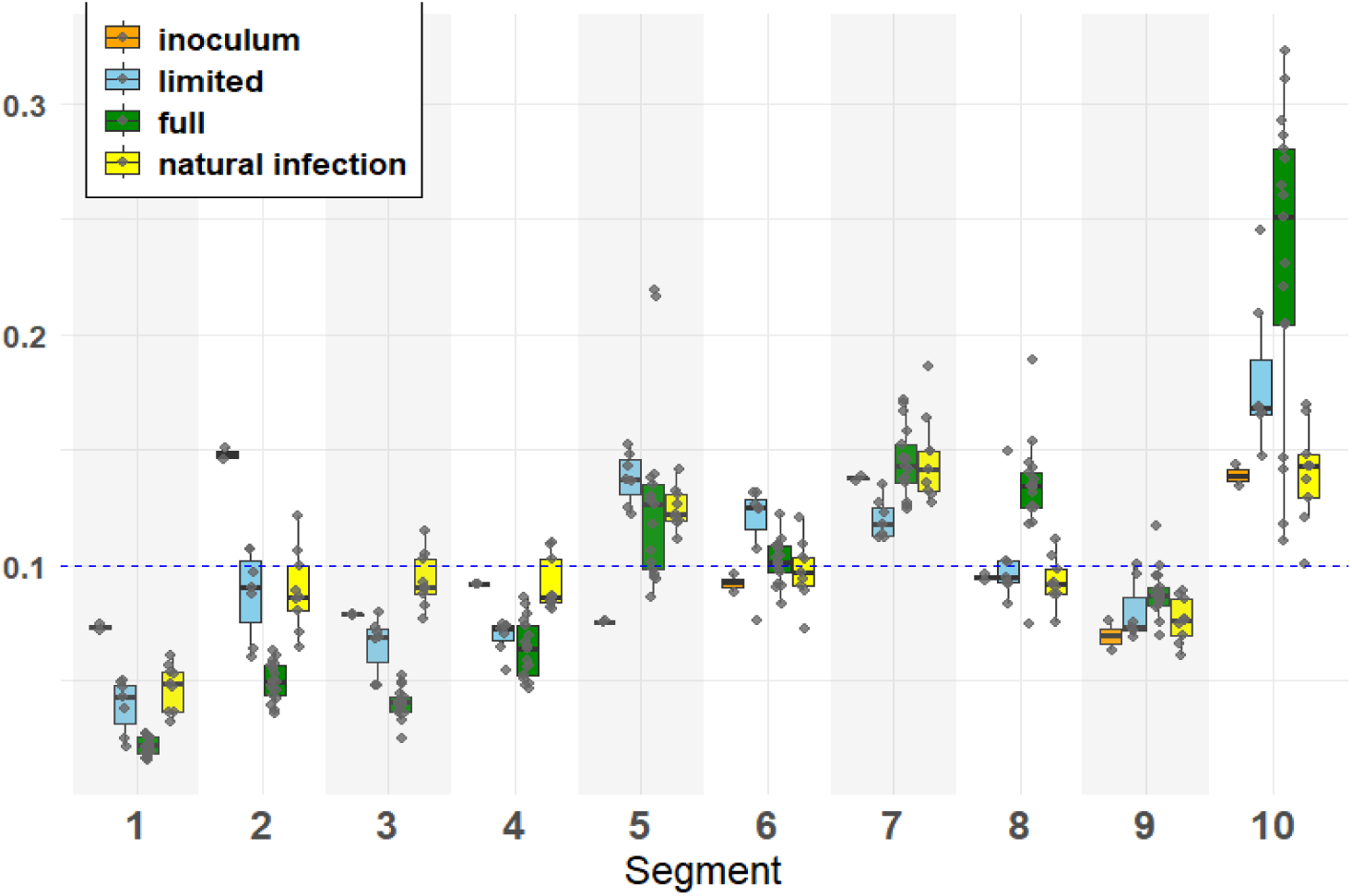
Frequencies of each BTV segment in four types of samples. Three types derive from the experimental infection of wild Culicoides imicola midges with BTV: (i) the inoculum and the midges with either (ii) a limited or (iii) a fully disseminated infection. The fourth sample type is naturally-infected midges from the field (data from [3]). Boxplots of each sample type are colored differently as shown in the legend. The horizontal dashed line indicates the expected frequency of the 10 segments at equimolarity. Due to the large number of different statistical analyses, graphical representations of p-values are not shown to improve visualization. Results of the comparisons between and within treatments are provided in the main text and in the supplementary material.

We also analyzed the correlation between dissemination level and viral load, which is another proxy of viral fitness. The viral load (*i.e*., the sum of the copies of all segments in a beheaded body) did not significantly differ between dissemination groups (Wilcoxon rank sum test, p-value = 0.18; Fig. S2). However, this observation remains tentative because we only included midges with relatively high viral loads to ensure a robust quantification of the GF. This choice could bias the analysis of the potential influence of the GF on viral load. Thus, we did not further explore this point given the difficulty of assessing both viral load and GF without a quantification bias.

Beyond the potential influence of GF on viral fitness, the experimental setup enabled us to address another unexplored question. We investigated whether the genome formula of BTV changes during transmission between different hosts. We thus compared segment frequencies between the inoculum and the midges. The mean frequency of most segments significantly differed between the inoculum and any of the dissemination groups (Student’s t test, Benjamini-Hochberg correction, p-values < 0.035 for most segment/dissemination combinations except for p-values > 0.05 for segment 7 in the full dissemination and segments 8 and 9 in the limited dissemination; Fig. 3, Tab. S7). Thus, the GF changed during transmission between hosts and did so toward specific values.

Finally, segment frequencies generally deviated from equimolarity in the same direction observed in natural infections (*i.e.,* above or below equimolarity). However, frequencies significantly differed between experimental and natural conditions for six segments (general linear model, p-value adjustment with rotation testing, p-values < 0.009 for segments 1, 2, 3, 4, 8 and 10; Fig. 3, Tab. S8).

### Deterministic processes participate to GFV generation in cell culture

The GF in the inoculum adjusted to specific values during midge infection in the previous experiment. Two main scenarios have been proposed to explain how GF changes [25]. The first scenario has two steps. First, early in host infection, different cells are infected by viral populations with different GFs due to random processes during cell infection. Thus, some cells would be infected with GFs closer to the optimal GF (*i.e.*, the set-point GF) than others. In a second step, viral populations with GFs closer to the set-point GF would have a higher fitness allowing them to become more abundant along infection. The alternative scenario postulates that differences in segment frequencies are due to differences in replication rates among segments. Thus, the first scenario implies random generation of GFs and subsequent selection, whereas the second scenario involves the deterministic generation of the set-point GF from the first round of infection.

To investigate how the set-point GF is generated, we analyzed GFs after a single round of replication in cell culture. This experimental design limits the influence of selection on the GF, favoring the detection of random processes. Moreover, the experiment was conducted in both vertebrate and insect cells to investigate putative differences between the two environments. Finally, for each cell line, two MOIs were used to analyze differences between infections caused by either a single virus particle (which is expected to carry all segments to yield a productive infection) or several particles (which allow complementation of incomplete genomes).

Under the assumption of strict random generation, the distributions of segment frequencies among infected cell cultures should overlap with those at equimolarity or in the inoculum. However, our observations did not follow this expectation (Fig. 4). Several segments had frequency distributions that significantly differed from equimolarity or the inoculum frequencies (Wilcoxon signed rank test, Benjamini-Hochberg correction, p-values < 0.05 for most segments but segment 9; Fig. 4 and Tab. S9). Those differences were found in all cell lines and MOI treatments (Fig. 4 and Tab. S9). Thus, segment frequencies significantly differed from equimolarity and from those in the inoculum after a single round of replication in cell culture.

**Figure 4.**
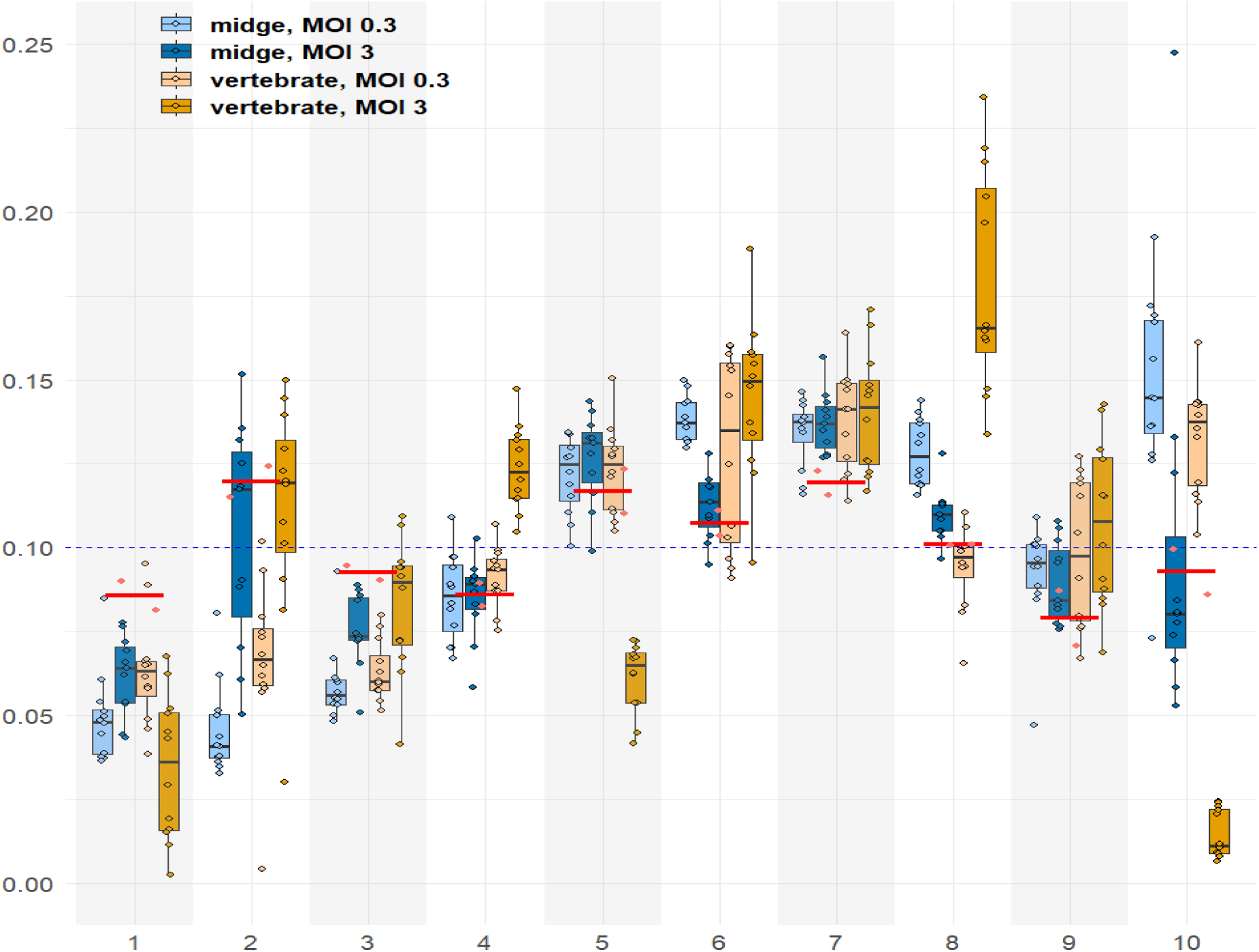
Frequencies of each BTV segment in cell culture infections (KC midge cells and MDBK vertebrate cells) at different MOIs (either 0.3 or 3). Boxplots of each combination cell line/MOI are colored differently as shown in the legend. The red bar and dots show the segment frequencies in the inoculum replicates. The horizontal dashed line indicates the expected frequency of the 10 segments at equimolarity. Due to the large number of different statistical analyses, graphical representations of p-values are not shown to improve visualization. Results of the statistical analyses between and within treatments are provided in the main text and in the supplementary material.

We then analyzed the influence of the cell line on the GF. As previously observed in natural populations, the frequencies of certain segments significantly differed between vertebrate and insect cells (general linear model per segment: frequency ∼ cell line, p-value adjustment with rotation testing, p-values < 0.03 for segments 4, 5 and 10; Fig. 4 and Table S10). Moreover, frequency distributions significantly differed between each cell line and natural populations from insect and vertebrate hosts (for each segment and host type, either midge or vertebrate: general linear model: frequency ∼ host, p-value adjustment with rotation testing, p-values < 0.05 for at least four segments; Fig. S3 and Tab. S11). Hence, although the cell lines differed from in-vivo conditions as previously observed [37], they represented different environments for the virus and led to specific genome formulas.

Finally, we analyzed how the MOI influences the GF in each cell line. We detected significant differences in the frequencies of certain segments between MOI treatments (general linear model per segment: frequency ∼ MOI, p-value adjustment with rotation testing, p-values < 0.02 for segments 2, 3, 6 and 8 in KC cells and for all segments except segments 6, 7 and 9 in MDBK cells; Fig. 4 and Tab. S12). There was no clear common trend in the influence of the MOI on the GF between the two cell lines. For example, increasing the MOI had a similar effect on the frequencies of segments 2, 3 and 10 in both cell lines, but had the opposite effect on segments 1 and 8 (Fig. 4).

## Discussion

This study explores three questions regarding host alternation in bluetongue virus (BTV). These questions are: (i) are host-specific mutations found in natural BTV populations?, (ii) does genome formula variation (GFV) influence viral fitness?, and (iii) what relative weight do random and deterministic processes have on GFV generation? Through the analysis of natural and experimental infections, we provide new insights into the evolutionary dynamics of BTV and the mechanisms that may facilitate its ability to cycle between vertebrate and arthropod hosts.

### Host Adaptation Through Mutation

Our study examined whether host-specific genetic changes take place during host alternation in BTV. Unlike previous studies [13,38], we analyzed field populations. No single-nucleotide polymorphism (SNP) was fixed across all samples from a given host. At most, the same fixed SNP was observed in half of the samples from a given host. Thus, host-specific mutations do not seem to be a major mechanism facilitating phenotype optimization at each host switch. Our results align with the few studies on wild populations of other arboviruses [14–17]. It is tempting to propose that host adaption through mutations might not often occur at each host switch in certain arboviruses. However, there are still too few studies on natural arbovirus populations to draw robust conclusions. Further research is needed to fully assess the potential role of host-specific mutations during host alternation with different arboviruses and in natural settings.

The absence of host-specific parallel mutations does not rule out an influence of the host species on BTV genetics. In fact, some of our findings could be due to such influence. For example, a larger number of non-parallel fixed mutations were detected in midges. This pattern may be due to severe bottlenecks during transmission from vertebrates to midges. That is, transmission would imply the random sampling of a small subset of the genetic diversity of infected vertebrates, potentially resulting in different genotypes fixed in each midge.

Our experimental design has several limitations that should be considered. First, due to the inherent properties of the sequencing approach, the extremities of the genomic segments were not fully sequenced. These regions could harbor host-specific mutations. Second, low-frequency mutations (frequency < 0.2) were not analyzed to limit false positives. Such mutations may contribute to phenotype optimization, for example via quasispecies dynamics [39]. Finally, we analyzed natural infections involving specific genetics of both BTV and its hosts, and in a specific environment (*i.e.* an epizooty of BTV serotype 4 in the Mediterranean region [3]). These infections may not reflect the behavior of all BTV strains in the nearly worldwide distribution of BTV.

### The Influence of GFV on Viral Fitness

We also investigated whether GFV influences viral fitness. To this end, we examined the relationship between the GF and the speed at which BTV disseminates in midges, which is a proxy of viral fitness. Our results revealed a correlation between specific genome formulas and the extent of viral dissemination within the midge. This correlation was observed following oral infection, the natural route of BTV transmission, and with wild midges. Thus, this study is one of the few showing a potential link between GFV and viral fitness [25,28,30,40]. However, the molecular mechanisms underlying this correlation cannot be clearly inferred from our data. For instance, many BTV segments exhibited changes in frequency, encoding both structural and non-structural proteins, which makes difficult to pinpoint a specific mechanism. Future research could analyze how the GF influences gene expression of both the host and BTV to identify potential mechanisms.

Beyond the link with fitness, this experiment allowed to study potential GF dynamics during between-host transmission. Previous observations of GFV in BTV populations could not address this question. In this study, the GF changed between the inoculum and the midge bodies and GFs tended to converge among midges. Thus, our results unveil the dynamic nature of the GF during host infection by BTV, as previously observed in multipartite viruses [25].

There are several limitations to our experimental design. First, we used viral detection in midge heads as a proxy for salivary gland infection and transmission probability, a common approach in BTV research [32–34]. However, we cannot rule out the possibility that some salivary glands were not fully contained within the dissected head tissue, which could lead to false negatives. Moreover, low viral loads prevented GF estimation in midge heads. Thus, we could not to assess potential GF dynamics during host colonization. Finally, differences were observed between GFs in experimental and natural infections. Despite using natural oral infection, blood meals spiked with BTV may not fully replicate the complexity of midge infection in nature.

### The Role of Random Processes in GFV Generation

The previous experiment and work on natural populations [3] have shown that the GF tends to converge towards specific values in a given host. These observations strongly suggest that GFV in BTV is not solely governed by random processes. Rather, selection and/or molecular mechanisms probably shape genome formulas toward specific values, known as set-point GFs. However, no data were previously available on the generation of the set-point GF in BTV. This paucity of data also pertains other viruses [41,42]. To address this question, we analyzed GF generation in cell culture experiments. The experiments were designed to limit selection pressures acting on BTV populations.

This experimental design unveiled that segment frequencies diverged from both the inoculum and equimolarity after a single round of replication. Additionally, frequencies converged toward different values in vertebrate and insect cells. These two lines of evidence strongly suggest that GFV is not generated only through stochastic processes. Rather, it appears that deterministic processes participate in GFV generation from the first cycle of infection. Moreover, these processes would act in a host-specific manner.

The molecular mechanisms that lead to the deterministic changes in segment frequencies are not known. In our opinion, two mechanisms may be involved: deletion of segment regions and encapsidation. However, these mechanisms are indistinguishable using our PCR quantification of segment copy numbers [3]. Therefore, different rates in deletion or encapsidation among segments could explain the observed differences in segment frequencies. However, current data on BTV encapsidation suggest that these phenomena may be rare. BTV encapsidation has been shown to result in the encapsidation of one copy of each segment via specific interactions among segment sequences [43]. This conservative encapsidation does not seem prone to generate defective particles. Nevertheless, most of our knowledge on BTV encapsidation comes from in-vitro or transfection experiments [23,24,43] and these experiments might not fully mimic natural infections. Moreover, several studies have identified defective particles with either deletions or non-conservative encapsidation in BTV populations [44,45]. Finally, viable deletion mutants have been generated using reverse genetics [46,47]. Overall, these studies suggest that BTV deletion mutants are sometimes encapsidated. Thus, BTV encapsidation may not always be fully conservative, as recently observed in other segmented viruses for which conservative encapsidation was thought to be the rule [48]. Future studies on the genomic content of BTV virions generated in natural hosts infections could help to explore potential mechanisms behind our results and BTV encapsidation in general.

## Conclusion

In summary, our findings suggest that BTV could optimize the phenotype at each host switch using strategies, if any is actually used, other than host-specific mutations. GFV seems to be an interesting candidate because it is well suited for phenotype optimization at each host switch. However, GFV must improve viral fitness to be considered as a strategy allowing phenotype optimization. Our study provides support for a link between GFV and viral fitness. Moreover, we show that GFV is not generated purely by random processes, supporting the hypothesis that mechanistic constraints shape genome formulas during cell infection. Future studies should further investigate the molecular mechanisms behind GFV and its potential role in optimizing BTV infection across hosts.

## Material and methods

### Illumina sequencing of field-collected BTV populations

#### Sampling and RNA extraction

Samples of natural BTV infections have been previously described in detail [3]. Briefly, cows, sheep and *Culicoides imicola* midges were sampled during a BTV-4 outbreak in two Mediterranean islands, Corsica (France) and Sardinia (Italy), between 2016 and 2017. Blood samples were obtained from symptomatic sheep in farms, and from cows during BTV routine testing in slaughterhouses. Cow samples were only collected in Corsica. Biting midges were collected using UV-light suction traps as previously described [49] but with a solution that limits RNase activity [50] in place of water in traps. *Culicoides imicola* females were identified on a cold plate, gathered in pools of 50 individuals, and stored at −20 °C.

RNA was extracted from the Corsican samples using the RNA Viral Extraction Kit (Macherey Nagel) following manufacturer’s instructions and using linear acrylamide (New England Biolabs) as carrier. Sardinian samples were extracted with the MagMAX CORE Nucleic Acid Purification kit (Applied Biosystems) following manufacturer’s instructions.

#### Sequencing protocol

We did not succeed in sequencing BTV genomes from field samples using an amplicon-based sequencing method (not shown). We thus developed a method to sequence field samples without the need for cell culture amplification. This method involves amplifying the BTV genome with a large number of primers situated throughout the genome. This approach is conceptually similar to previous methods designed to sequence viruses in samples with low viral loads or highly degraded [51]. One specificity of our method is the use of primers with random hexamers at their 3’ extremities to generate double-stranded DNA (dsDNA) from cDNA. This strategy should avoid a major drawback of virus sequencing methods based on only virus-specific primers [51], that is the limited amplification mutants diverging from the sequences of virus-specific primers. Another specificity that distinguishes our method is the library construction step. We have adapted an in-house library preparation, thus reducing the cost per library [52]. A detailed protocol is provided in Annex 1.

Briefly, a reverse transcriptase reaction was used to generate cDNAs from BTV segments with the RevertAid First Strand cDNA synthesis kit (Fisher Scientific) and using 186 BTV-specific primers (Table S13). The primers were designed with Geneious (version R10) to be distributed throughout the BTV-4 genome at approximately a 100-bp interval. The final set of primers was selected after validating adequate genome coverage during sequencing of nine samples from our collection (see above) in an Illumina MiSeq sequencer (250-bp paired-end mode; not shown). The cDNA then was converted to double-stranded DNA (dsDNA) using the Klenow fragment (Fisher Scientific) and a combination of primers either targeting the 5’ end of each segment (primers with the term 5SPE-TAGE at the end of their name in Table S13) or containing a random hexamer at their 3’ extremities. Both BTV-specific and random primers have the same sequence in their 5’ ends (i.e. 454-E primer sequence, Annex 1). The dsDNA fragments were amplified in a PCR reaction using the 454-E primer. Adaptor sequences were added in a subsequent PCR reaction (Phusion High-Fidelity DNA Polymerase kit, Fisher Scientific) with primers bearing either Illumina P5 or P7 adaptor sequences (Annex 1). Library sizing targeted 600-bp amplicons and was done using the AmpurXP magnetic bead capture system (Agencourt; 0.65X bead volume) following manufacturer’s instructions. Library size was validated using capillary electrophoresis (Agilent 2100 Bioanalyzer, Agilent Technologies). Library quantification was done with the Library Quantification kit (Takara Bio) according to the manufacturer’s protocol.

Twenty-one libraries were generated from field samples, plus two negative control libraires (extraction and amplification controls). The libraries were sequenced in full run of a HiSeq2500 (Illumina, 250-bp paired-end reads) by Macrogen (South Korea). All reads have been deposited in the Sequence Read Archive (SRA) database under the Bioproject accession PRJNA1349788.

#### Bioinformatic analysis

Raw reads were analyzed for nucleotide variations using the Varhap pipeline [53]. Varhap is freely available at https://github.com/loire/Varhap. Varhap calls for modules available in SAMtools [54], including alignment of reads with BWA [55] and variant calling with freebayes [56]. Variant Call Format (VCF) files summarized the exact proportion of each variant detected. Variants were determined by applying a filter on raw data. The filter used to detect variants in this study mainly was (vcffilter -f “QA/AO > 10 & AO/RO > 0.01 & DP > 50 & RPL > 2 & RPR > 2 & SAF > 2 & SAR > 2”) where QA represents the addition of base? “A” qualities, A0 (A for Alternative) represents the number of reads on the alternative base, R0 (R for reference) represents the number of reads on the reference base, DP represents the depth, RPR and RPL reads “balanced” to each side (left and right) of the site. SAF/SAR represent the number of reads on the forward and reverse sequences.

In a first step, consensus sequences were derived from a BTV-4 strain sampled along with the samples used in this study (GenBank accessions: KY654328.1-KY654328.10). Analysis of SNPs showed 35 mutations fixed or near fixation in at least ten samples and in at least two samples of each host. These differences could be due to cell culture passages of the BTV-4 strain used as consensus. Thus, we generated a new consensus including the mutations observed at a frequency above 0.9 and in at least ten samples. The sequences of the primers targeting segment extremities were removed (approximately 27-29 pb) from the analysis. The total genome length included in the analysis was 18571 pb.

### Quantification of BTV segments with reverse transcription – real-time PCR (RT-qPCR)

The RT-qPCR approach has previously been described [3]. This method uses ten qPCR designs (one per segment) to quantify only genomic segments. Viral transcripts are not amplified because the RT step targets the negative strand of the segments and this strand, which is only generated within fully-formed capsids [57]. Briefly, the approach involves a two-step RT-qPCR for each segment. Primers and cycle parameters are provided in Tables S14 and Annex 2, respectively. A control sample was included in all PCR plates to ensure the ten qPCRs were accurately calibrated. The control sample contained a plasmid harboring a concatenate of the ten regions targeted by the qPCRs [3]. Thus, the control sample has the same number of copies of the ten qPCR targets and thus allows to calibrate all qPCRs using the same control. When the plasmid was used as template for the ten qPCRs, the copy numbers were nearly equimolar (i.d., frequencies = 0.1). For example, maximum and minimum frequencies of any segment were between 0.106-0.096 and 0.247-0.003 for the control and the cell culture infections, respectively. All samples were analyzed in triplicate.

We analyzed the potential influence of primer-independent cDNA synthesis in RT-qPCR reactions as previously described [3]. This phenomenon occurs when nucleic-acid molecules (*e.g*., tRNA) or RNA secondary structures act as primers in the RT reaction [58]. We have previously shown that this phenomenon had a limited influence on the quantification of BTV segments [3]. Copy number did not significantly differ between datasets corrected or not for primer-independent cDNA synthesis (Wilcoxon, p-values > 0.058). All statistical analyses were done on the dataset corrected for copy number overestimation due to primer-independent cDNA synthesis.

### Experimental infection of wild *Culicoides* biting midges

The capture of biting midges and their experimental infection have previously been described in detail [34]. A description of each step is presented below.

#### Midge trapping

Biting midges were captured on a farm in Sardinia (Italy; Bari Sardo municipality) in 2016. UV-light suction traps were set up for an overnight capture. Captures were carried out for two to three consecutive nights to collect a sufficient number of midges. Midges were collected each morning and kept in cardboard boxes for about two days. During this period, the midges were fed with a 10% sucrose solution and kept at 25 °C. Then, the midges were starved for one day before experimental infection.

#### Experimental infection of *Culicoides imicola* females

Virus-spiked blood meals were prepared through diluting BTV suspensions (10^5.3^ – 10^6.5^ PFU/mL) in sheep blood (1:3 dilution). The BTV strain used was BTV-4 ITA, strain 2014TE31172 (REF). This strain was isolated from the blood of a sheep in the Apulia region (Italy) in 2014. The strain was passed once onto the KC cell line (derived from *Culicoides sonorensis* midges) and three times onto VERO cells before use [59]. Midges were fed on blood meals using a Hemotek feeding system for 45 minutes. Then, midges were anesthetized and sorted by species. Only *Culicoides imicola* females were used in the following steps. Fully-engorged females were kept at 25 °C, with ad libitum access to a 10% sucrose solution. Ten days later, the surviving midges (n = 258) were frozen on dry ice. Their heads and bodies were separated and stored at −80 °C.

#### Nucleic acid extraction and BTV screening in heads and beheaded bodies

Each head or beheaded body was homogenized in PBS 1X using a TissueLyser II (Qiagen). Nucleic acids in heads were extracted using the Biosprint^®^ 96 kit (Qiagen). Nucleic acids in beheaded bodies were extracted with the MagMAX Core Nucleic Acid Purification kit (Applied Biosystems) with linear acrylamide as carrier.

A routine diagnostic RT-PCR for BTV was used to detect BTV in midge heads [60]. Additonaly, a serotype-specific RT-PCR (VetMAX European BTV Typing Kit, Thermo Fisher Scientific) was also performed on the heads to distinguish between infections by the strain used in the experimental infection and other BTV serotypes potentially circulating in the region (serotypes 2, 8 and 16). Only individuals that tested negative for natural BTV infections were used.

Segment abundance was quantified in the beheaded bodies of 30 females with BTV-4 positive heads. RT-qPCRs targeting each segment (see above) were used to quantify segment copy numbers. We defined a Ct threshold of 31 to guarantee robust quantification of segment copies. Seventeen females showed Ct values below 31 for all segments. These females were considered the group with a full virus dissemination. Then, we screened the beheaded bodies of 40 females with BTV-negative heads. Seven females showed Ct values below 31 for all segments. These seven females formed the group with a limited virus dissemination.

### Cell culture infection

#### Virus

The FCO-4_K1B2 strain was used for cell culture infections. This strain is derives from a BTV-4 isolate obtained from sheep blood collected in Corsica (France), in 2016 [59]. This isolate was obtained after inoculation of KC cells with sheep blood. The isolate was passage twice in BSR cells (a clone of BHK-21 cell line, from hamster kidney cells) to obtain the FCO-4_K1B2 strain. All experiments were carried out with the stock generated during the second passage in BSR cells.

The 50% tissue culture infectious dose (TCID50) of the FCO-4_K1B2 stock was determined using standard endpoint dilution on Madin-Darby Bovine Kidney epithelial (MBDK) and *Ovis aries* testis (OA3.Ts) cells, as described [61]. The multiplicities of infection (MOIs) were estimated using plaque-forming units per ml (PFUs). The latter were calculated multiplying TCID50 values by 0.69, as previously described [62]. Then, the appropriate PFUs per cell were estimated for each MOI treatment.

#### Cell culture infection and nucleic acid extraction

KC and MDBK cells were inoculated with the FCO-4_K1B2 strain at two MOIs: 0.3 and 3. Titration on KC cells was not possible because BTV does not lead to plaques in KC cells. Thus, the MOIs in KC cells were based on a titration in OA3.Ts cells. The OA3.Ts cells are more susceptible to BTV infection than MDBK cells (*i.e.,* the same virus solution titers higher in OA3.Ts cells than in MDBK cells). Therefore, titration in OA3.Ts cells was chosen to maximize the chances of obtaining an MOI below one when inoculating KC cells with the lowest MOI. Using this approach, virus concentration in the inoculums for KC cells was half of that used for MDBK cells.

For each MOI treatment and cell line, cells at confluence were washed three times with 1X PBS buffer, inoculated with a virus suspension for two hours, and washed again with 1X PBS buffer before the corresponding medium was added. To obtain a similar number of infected cells between treatments, the number of initial cells in the wells differed between MOI treatments for each cell line (cell numbers for KC line = 8.0E5 and 2.0E5, cell numbers for KC line = 3.2E5 and 8.0E4). This approach was intended to minimize biases in segment quantification due to unequal sizes of viral populations between treatments. The cells were aliquoted (150 µl/aliquot) and frozen at −80°C twelve hours after inoculation. This time period has been shown to allow for a single replication cycle [23]. Infections for each MOI/cell line treatment were done in triplicate and in parallel. The whole experiment was repeated twice on different days.

An aliquot of frozen cells (150 µl) was used for each extraction. Nucleic-acid extraction was done using the MagMAX Core Nucleic Acid Purification kit (Applied Biosystems) following manufacturer’s instruction and using linear acrylamide as carrier.

### Statistical analyses

Statistical tests were carried out with the R software (R Development Core Team, 2011, version 4.3.0). Distance of segment frequencies to equimolarity were calculated as 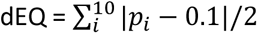 according to Manly (eq. 5.7 p. 68) [63], where i is the segment, p is the relative frequency of the segment in the sample and 0.1 is the relative frequency at equimolarity.

## Ethics statement

The sampling was performed by competent public bodies in charge of bluetongue surveillance. More precisely, in France, sample collection was carried out by the French National Reference laboratory for bluetongue (Agence Nationale pour la sécurité alimentaire), in collaboration with the local Veterinary services. In Italy, collections were carried out by the Italian National Reference laboratory for bluetongue (Istituto Zooprofilattico Sperimentale dell’Abruzzo e del Molise “G. Caporale”), in collaboration with the network of the Italian Istituti Zooprofilattici Sperimentali and the local Veterinary Services.

## Acknowledgments

This work was supported by research grants CuliOme (ANIHWA, ERA-Net) and PALE-Blu (H2020) from European Union. This work was also funded by the Direction générale de l’alimentation from the French Ministry of Agriculture and Food. We are grateful to the Directions départementales de la cohésion sociale et de la protection des populations from South Corsica and High Corsica, and to the Groupement Défense Sanitaire Bétail from High Corsica, for their support in collecting biting midges.

## Supporting information captions

### Supplementary figures

**Figure S1.**
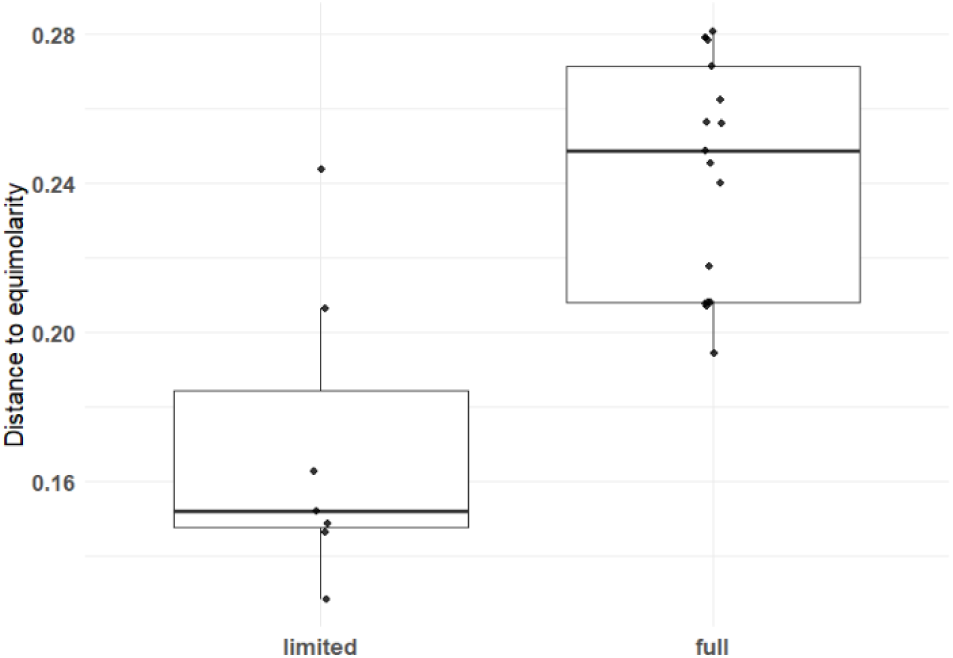
Distance to equimolarity of segment frequencies in the two dissemination groups, the full and limited-dissemination groups.

**Figure S2.**
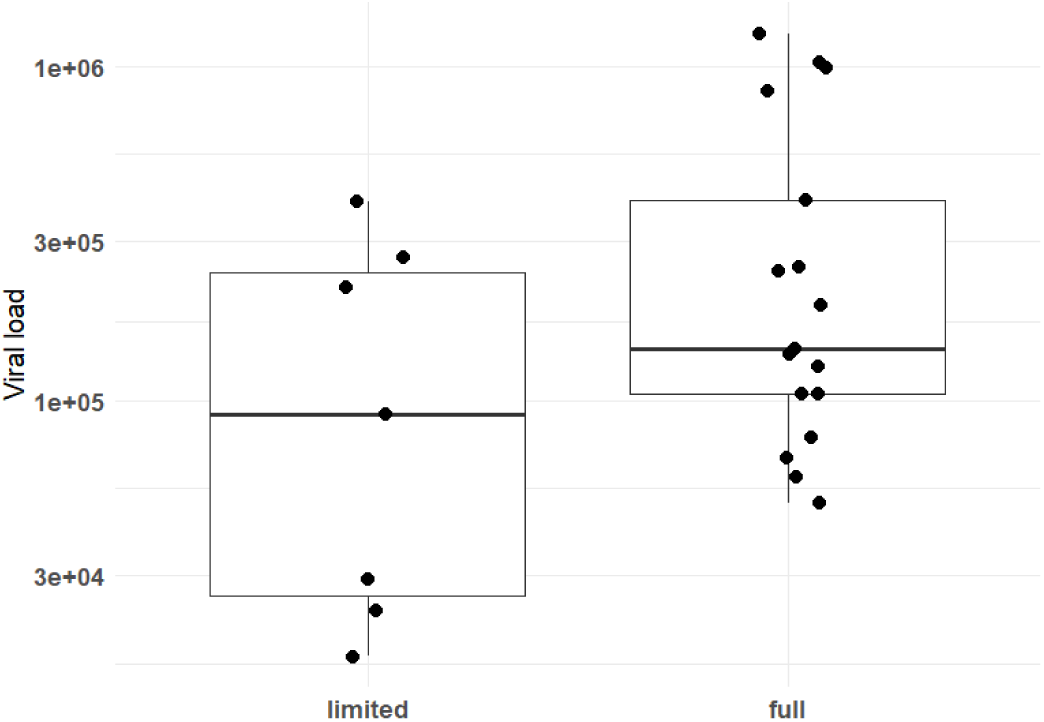
Viral loads in the limited and full dissemination groups.

**Figure S3.**
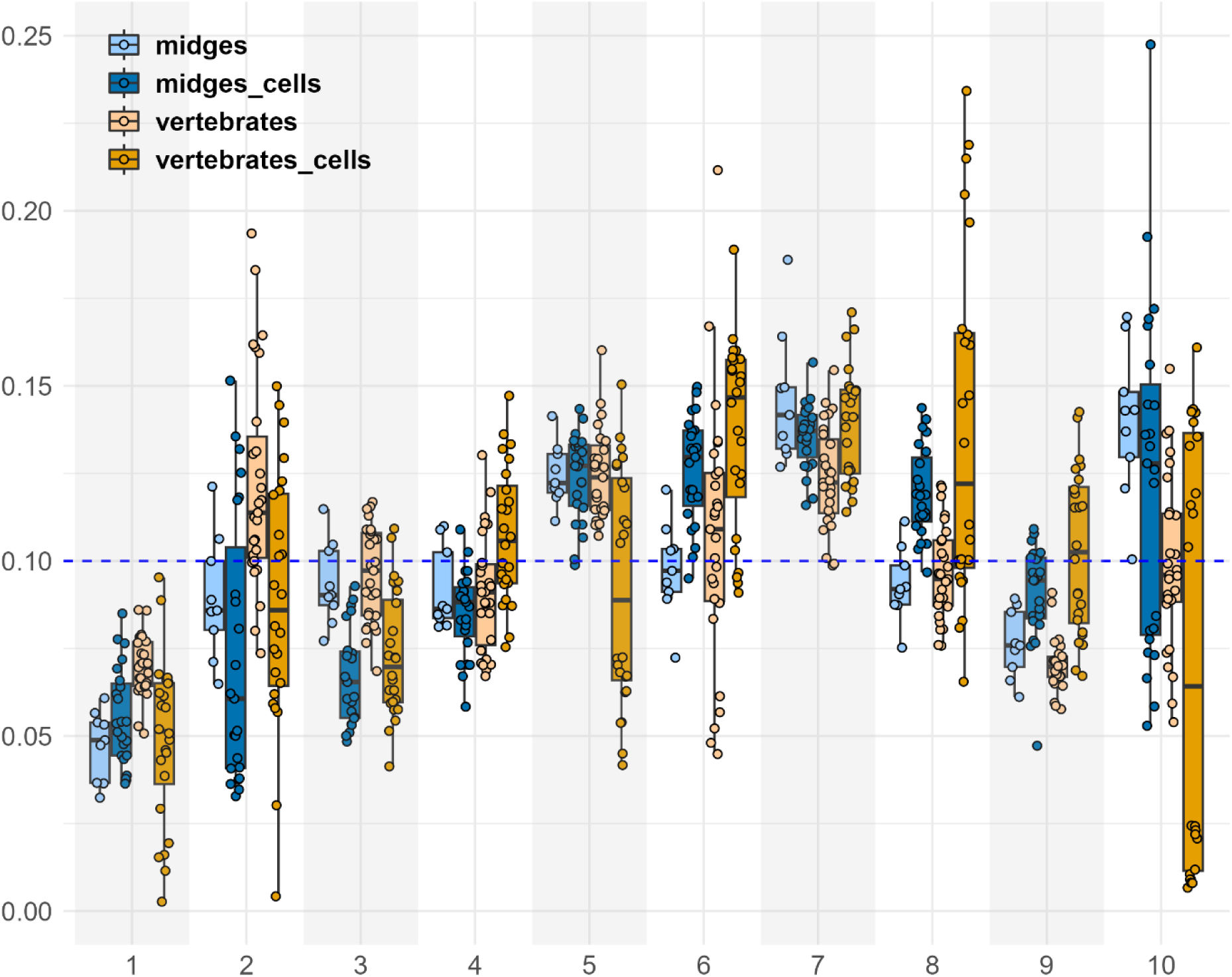
Frequencies of each BTV segment in cell culture infections and in naturally-infected hosts. Boxplots of each host type are colored differently as shown in the legend. Host type names ending by “_cells” identify cell culture samples (midges_cells = KC cells, vertebrates_cells: MDBK cells). The host type “vertebrates” includes samples from both cows and sheep. Due to the large number of comparisons, graphical representations of p-values are not shown to improve visualization (see Tab. S7). Segments showing significant differences for each host type: 3, 6, 8 et 9 for arthropod hosts, and all segments but segment 10 for vertebrate hosts (Tab. S7).

### Supplementary tables

Table S1. Single nucleotide polymorphisms (SNPs) along the genome of Bluetongue virus and among hosts. SEG: segment, POS: position of the SNP on a given segment, SEG_POS: concatenation of the segment and the SNP position on the segment, REF: nucleotide in the reference sequence, ALT: nucleotide in the sample, FREQ: SNP frequency, HOST: host species.

Table S2. Fixed single nucleotide polymorphisms (SNPs) shared by two samples of the same host.. SEG: segment, POS: position of the SNP on a given segment, HOST: host species, REF: nucleotide in the reference sequence, ALT: nucleotide in the sample, REF_codon: codon in the reference sequence, REF_AA: aminoacid in the reference sequence, ALT_codon: codon in the sample sequence, ALT_AA: aminoacid in the sample sequence, Non-synonymous: “1” stands for SNPs leading to a non-synonymous change.

Table S3. Four non-fixed SNPs found in several samples of the same host. SEG: segment, POS: position of the SNP on a given segment, HOST: host species, FREQ: SNP frequency, REF: nucleotide in the reference sequence, ALT: nucleotide in the sample, REF_codon: codon in the reference sequence, REF_AA: aminoacid in the reference sequence, ALT_codon: codon in the sample sequence, ALT_AA: aminoacid in the sample sequence, Non-synonymous: “1” stands for SNPs leading to a non-synonymous change.

Table S4. Comparison of segment frequencies with equimolarity in laboratory-infected midges.

Table S5. Comparison of frequencies between segments in laboratory-infected midges.

Table S6. Comparison of segment frequencies between the two groups of laboratory-infected midges with either a full or a limited dissemination of the viral infection.

Table S7. Comparison of the mean frequencies of each segment between the inoculum and each the dissemination level.

Table S8. Comparison of segment frequencies between BTV populations from either field or experimental infections of biting midges

Table S9. Comparison of segment frequencies with equimolarity and the inoculum in cell culture infections. The two cell lines are the KC insect cell line and the MDBK vertebrate cell line.

Table S10. Comparison of segment frequencies between BTV populations in the KC insect cell line and the MDBK vertebrate cell line.

Table S11. Comparison of segment frequencies between BTV populations in cell lines and field populations in the same host type (i.e., KC insect cell line versus field-caught midges and sheep, and MDBK vertebrate cell line versus sheep and cows sampled during an epizooty).

Table S12. Comparison of segment frequencies between BTV populations in cell culture infections at two multiplicities of infection (MOIs) either in the KC insect cell line or the MDBK vertebrate cell line. The MOI values were 0.3 and 3.

Table S13. Primers used to sequence populations of Bluetongue virus in field samples.

Table S14. Primers used for the quantification of the negative genomic strand of BTV-4. The “Forward” primers were used in separate mixes for reverse transcription at a final concentration of 2 µM.

Table S15. Reaction and cycling conditions for the RT-qPCRs. A) Reagents and their volumes. Reactions were prepared separately for each segment. An additional reaction was prepared for each sample in which the primer was replaced with water to test for primer-independent cDNA amplification. B) temperature and length of each RT-qPCR step.

### Supplementary annexes

Annex 1. Protocol for the preparation of libraries for Illumina sequencing used to characterize the population genetics of Bluetongue virus in field samples.

Annex 2. Protocol for the quantification of genomic segments of Bluentongue virus with real time PCR.

